# Brain wiring and supragranular-enriched genes linked to protracted human frontal cortex development

**DOI:** 10.1101/746248

**Authors:** Jasmine P. Hendy, Emi Takahashi, Andre J. van der Kouwe, Christine J. Charvet

**Affiliations:** Department of Biology, Delaware State University, DE, 19901, USA; Division of Newborn Medicine, Department of Medicine, Boston Children’s Hospital, Harvard Medical School, Boston, MA, 02215, USA; Fetal-Neonatal Brain Imaging and Developmental Science Center, Boston Children’s Hospital, Harvard Medical School, Boston, MA, 02215, USA; Athinoula A. Martinos Center for Biomedical Imaging, Massachusetts General Hospital, Harvard Medical School, Charlestown, MA, 02129, USA; Center for Neuroscience, Department of Psychology, Delaware State University, DE, 19901, USA

**Keywords:** Cortex, development, evolution, primates, humans, supragranular-enriched genes

## Abstract

The human frontal cortex is unusually large compared with many other species. The expansion of the human frontal cortex is accompanied by both connectivity and transcriptional changes. Yet, the developmental origins generating variation in frontal cortex circuitry across species remain unresolved. Nineteen genes, which encode filaments, synapse, and voltage-gated channels (e.g., *NEFH*, *SYT2*, *VAMP1*) are especially enriched in the supragranular layers of the cerebral cortex in humans relative to mice. The increased expression of these genes suggests enhanced cortico-cortical projections emerging from layer III in humans. We confirm that the expression of these supragranular-enriched genes is preferentially expressed in frontal cortex layer III in humans relative to mice. We demonstrate a concomitant expansion in cortico-cortical pathways projecting within the frontal cortex white matter in humans with diffusion MR tractography. To identify developmental sources of such variation, we compare frontal cortical white matter growth and developmental trajectories of transcriptional profiles of supragranular-enriched genes in humans and mice. We also use temporal changes in gene expression during postnatal development to control for variation in developmental schedules across species. The growth of the frontal cortex white matter and transcriptional profiles of supragranular genes are both protracted in humans relative to the timing of other transformations. These findings demonstrate that an expansion of projections emerging from the human frontal cortex is achieved by extending the duration of cortical circuitry development. Integrating RNA sequencing with neuroimaging level phenotypes is an effective strategy to assess deviations in developmental programs leading to variation in connections across species.

## Introduction

The human frontal cortex is unusually large compared with that of many other species such as rodents, though the claim that the frontal, and in particular the prefrontal cortex, is unusually large in humans relative to apes remains controversial (Sherwood and Smaers, 2013; Semendeferi et al., 2002; Barton and Venditti, 2013; Carlén, 2017; Donahue et al., 2018). The expansion of the frontal cortex in humans versus rodents reflects modifications to its underlying molecular architecture and patterns of connectivity.

Patterns of gene expression across the layers of the cerebral cortex are generally conserved across primates and rodents, but there are some notable differences across species. A study profiling ~1,000 genes found that 79% have conserved pattern of expression across layers in humans and in mice (Zeng et al., 2012), but that 21% exhibited species-specific variation. Of particular interest, 19 genes (e.g., *NEFH*, *SYT2*, *VAMP1*) are especially enriched in the supragranular layers (i.e., layer III) of the cerebral cortex in humans relative to mice (Zeng et al., 2012; Krienen et al., 2016). These genes encode filaments (e.g., *NEFH*), synapse (e.g., *VAMP1*), and voltage-gated channels (e.g., *SCN4B*). The differential expression of these genes across cortical layers suggests major modifications to cross-cortical connectivity patterns between humans and mice. Because supragranular layer pyramidal neurons (i.e., layer III neurons) generally project within the white matter to target various cortical areas, the increased expression of genes encoding filaments and voltage-gated channels in layer III of humans suggest enhanced cortico-cortical projections coursing through the white matter in humans relative to mice (Gilbert and Kelly, 1975; Kennedy and Bullier, 1985).

The notion that humans as other primates possess enhanced cross-cortical projections compared with other mammals is consistent with the observation that primates possess an expansion of upper layer (i.e., layer II-IV) neurons (Charvet et al., 2017), some of which form long-range cortico-cortical projections (Charvet et al., 2019a). The expansion of layer III neuron numbers in the human frontal cortex is also evident from single cell sequencing studies, which show that excitatory layer III neurons represent ~46.5% of excitatory frontal cortex neurons in humans, but only ~24% in mice (Fig. 1; Luo et al., 2017). Thus, major differences in cell type organization across the depth of the cortex are associated with modifications to frontal cortical circuitry in humans.

**Figure 1.**
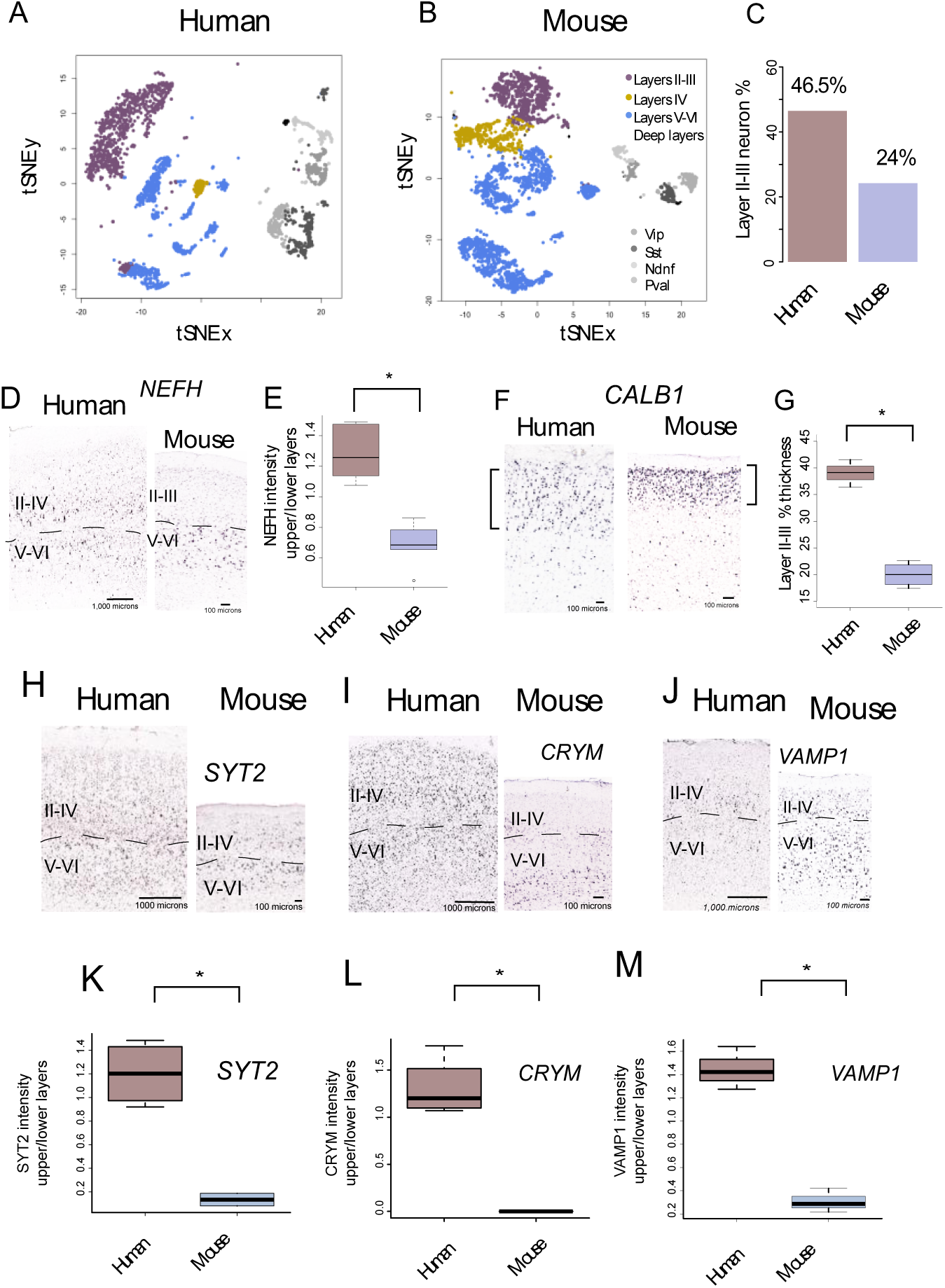
Single-cell methylome sequencing through the frontal cortex of a human(A) and a mouse (B) show that the human frontal cortex contains relatively more excitatory layer III neurons (C) compared with mice. Excitatory layer III neurons consist of 46.5% of frontal cortical excitatory neurons whereas excitatory layer III neurons only consist of 24% in mice. (D) *NEFH* expression is relatively and significantly increased in upper layer of humans compared with mice (E). Moreover, the thickness of layer III as assessed by *CALB1* is significantly expanded in humans compared with mice (G). These data together suggest major modifications to long-range cortical pathways emerging from layer III of the frontal cortex in humans compared with mice. Of the supragranular enriched genes, *SYT2* (H), *CRYM* (I), *VAMP1* (J) expression are significantly increased in upper versus lower layers in humans relative to mice (K-M). Sections of *NEFH, CRYM, SYT2, and VAMP1* mRNA expression are from the Allen brain atlas 2010 Allen Institute for Brain Science. Allen Human Brain Atlas. Data for humans are available from: human.brain-map.org.

We compare the frontal cortex of humans and mice because of their obvious differences in structural organization, which we use to develop a framework to integrate neuroimaging with molecular level phenotypes. Integrating across scales of organization from neuroimaging to molecular level phenotypes is a challenging enterprise to identify evolutionary changes in connections and their underlying developmental programs. Previous methods such as tract-tracers, diffusion MR tractography or carbocyanine dyes suffer from a number of limitations such that the study of evolutionary changes in connectivity patterns has largely remain untackled. Accordingly, this study is as much a proof of concept with which to link multiple scales of organization as it is a study to identify deviations in developmental programs giving rise to the emergence of frontal cortical circuitry in humans (Fornito et al., 2019). We select humans and mice as we can build from prior work to identify deviations in developmental programs, which is not as extensive in apes.

Human development is unusually long compared with that of mice and many other species (Clancy et al., 2001; Workman et al., 2013). Identifying deviations in the timing of developmental programs requires controlling for overall changes in developmental schedules across species (Charvet et al., 2017b) because developmental processes take longer in humans compared with many other species. Previous work tackling evolutionary changes in developmental programs between humans and other species has employed various strategies to compare developmental schedules across species. Those include the use of actual age, birth, or puberty as a basis with which to compare developmental schedules across species (Finlay and Workman, 2013; Sakai et al., 2017; Hawkes and Finlay, 2018; Charvet and Finlay, 2018). Relying on a few developmental transformations such as birth can be problematic. For example, birth does not align with the timing of most other neurodevelopmental events, and should therefore not be used as a basis with which to find corresponding ages across species (Finlay and Darlington, 1995; Clancy et al., 2007; Hawkes and Finlay, 2018; Charvet and Finlay, 2018). Rather than relying on the timing of a select few developmental transformations, we use the translating time model, which relies on the timing of 271 developmental events gathered across 19 mammalian species to find corresponding ages between humans and mice to infer deviations in developmental programs generating variation across the two species (Workman et al., 2013). Because the translating time model extends up to two years of age in humans and their equivalent in other species, we use temporal changes in gene expression from RNA sequencing from the frontal cortex of humans and mice at different postnatal ages to find corresponding time points up to 30 years of age in humans and their equivalent in mice.

We compare transcriptional profiles of supragranular-enriched genes from the frontal cortex of humans versus mice in adulthood and in development because they encode filaments (e.g., *NEFH*) and voltage-gated channels which are linked to long-range cortico-cortical pathways (Hof et al., 1995; Zeng et al., 2012; Krienen et al., 2014; Nguyen et al., 2017). We also compare the developmental trajectories in the growth of the frontal cortex white matter between humans and mice because the white matter houses projections coursing to and from the frontal cortex, which permits testing whether there are deviations in the timing of projection development coursing within, to or from the frontal cortex across these two species. Our findings demonstrate that protracted developmental programs associated with the development of long-range projections accounts for the expansion of frontal cortex cross-cortical circuitry in humans.

## Results

We first confirm that layer III is expanded in the frontal cortex of humans compared with mice. We then identify the developmental basis of such variation in frontal cortical connections between humans and mice. We focus on temporal changes in supragranular gene expression, as well as the growth of the frontal cortex white matter in humans and in mice to identify deviations in developmental programs accounting for the expansion of the human frontal cortex.

### Expansion of layer III and long-range projections in humans in adulthood

We perform a number of analyses to identify differences in layer III neuron organization in the frontal cortex of humans versus mice in adulthood. Single cell methylome sequencing from the frontal cortex of humans and mice demonstrate that the relative number of excitatory layer III versus excitatory layer IV-VI neurons in humans (layer III neurons: 873, layer IV-VI neurons: 999) is greater than expected by chance compared with mice (excitatory layer III neurons: 690; excitatory layer IV-VI neurons: 2,160; *χ*^2^=256.56, p<0.05; Fig. 1A-B). That is, 46.5% of excitatory frontal cortical neurons are located in layer III in humans but only 24% of excitatory layer III are observed in mice (Fig 1C; (Luo et al., 2017). If layer III neurons projecting over long distance are indeed expanded in the frontal cortex of humans, we would expect genes expressed by large neurons in supragranular layers to be increased in humans relative to mice. *NEFH* is typically expressed by large neurons, which often project over long distances. In the frontal cortex, we previously showed that *NEFH* expression in upper versus lower layers in humans is significantly greater than in mice (n=6 humans, n=6 mice, t=6.5318; p<0.05; Fig. 1D-E), further demonstrating an expansion of long-range projecting neurons in the frontal cortex of humans relative to mice. We also compare the relative expression of *CALB1* across the depth of the cortex in these two species because it is a marker for layer II-III (see Fig. S1; Nieto et al., 2004). The relative thickness of layer III as assessed from *CALB1* expression (a marker for layer II-III neurons; See Fig. S1) is significantly expanded in the frontal cortex of humans compared with mice (t=-10.185, p<0.05, n=4 mice 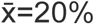, 3 humans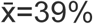, Fig. 1F-G), as is the relative thickness of layer II-III as defined cytoarchitecturally (Fig. S1). Together, these findings suggest an expansion and major modifications to layer III neurons in the human frontal cortex compared with mice.

Concomitant with the expansion of layer III neurons in humans versus mice, the human frontal cortex also possesses a suite of genes preferentially expressed in upper layers of the visual cortex (i,e., layer III), which encode filaments synapse and voltage-gated channels (Zeng et al., 2012). To confirm the supragranular enrichment in upper layers in the frontal cortex of humans relative to mice, we quantified the relative intensity of expression of some supragranular-enriched genes *(i.e., SYT2*, *CRYM*, and *VAMP1)* in upper versus lower layers across the frontal cortex of humans and mice (Fig. 1H-J). This is because prior work focused on comparing the expression of these genes focused on the primary visual cortex (Zeng et al., 2012), and it is not clear whether species differences in the expression of these supragranular-enriched genes are also evident for the frontal cortex. In the frontal cortex, *SYT2, CRYM*, and *VAMP1* are significantly preferentially expressed in upper layers in humans compared with mice (*SYT2*: t= −7.3234, p<0.05, n=4 humans 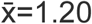, 2 mice 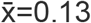; *CRYM*, t= −8.3528, p<0.05, n=4 humans 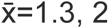, 2 mice 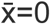; *VAMP1*, t= −9.3108, p<0.05, n=3 humans 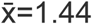, n=3 mice 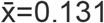; Fig. 1K-M). These findings demonstrate that humans deviate from mice in possessing major modifications in gene expression within layer III, which may be a molecular signature underlying major modifications to long-range corticocortical projections emerging from layer III neurons in the frontal cortex.

Diffusion MR tractography permits visualizing pathways across the entire brain of different species, and our quantitative comparisons across layer III of the human and mouse frontal cortex suggests enhanced cortico-cortical pathways in humans. We therefore test for the expansion of cortico-cortical pathways in the frontal cortex of humans with the use of high angular resolution diffusion MR tractography. An ROI placed through the white matter of the frontal cortex shows a number of cortico-cortical pathways such as the arcuate fasciculus and uncinate fasciculus in humans (Fig. 2A), which are not observed in mice (Fig. 2B). Horizontal slice filters used to capture fibers coursing to and from the frontal cortex in mice only identify the cingulate bundle, the corpus callosum Fig. 2B), and cortico-subcortical pathways (i.e., fibers coursing across the dorsal to ventral direction). We do not observe many cortico-cortical pathways coursing through the frontal cortex in mice. This is in contrast with humans where horizontal slice filters through the human brain show a number of long-range projections and U fibers coursing through the frontal cortex of humans (Fig. 2C-D). These qualitative observations from diffusion MR tractography show-case major differences in long-range cortico-cortical pathways emerging to or from the frontal cortex across the two species.

**Figure 2.**
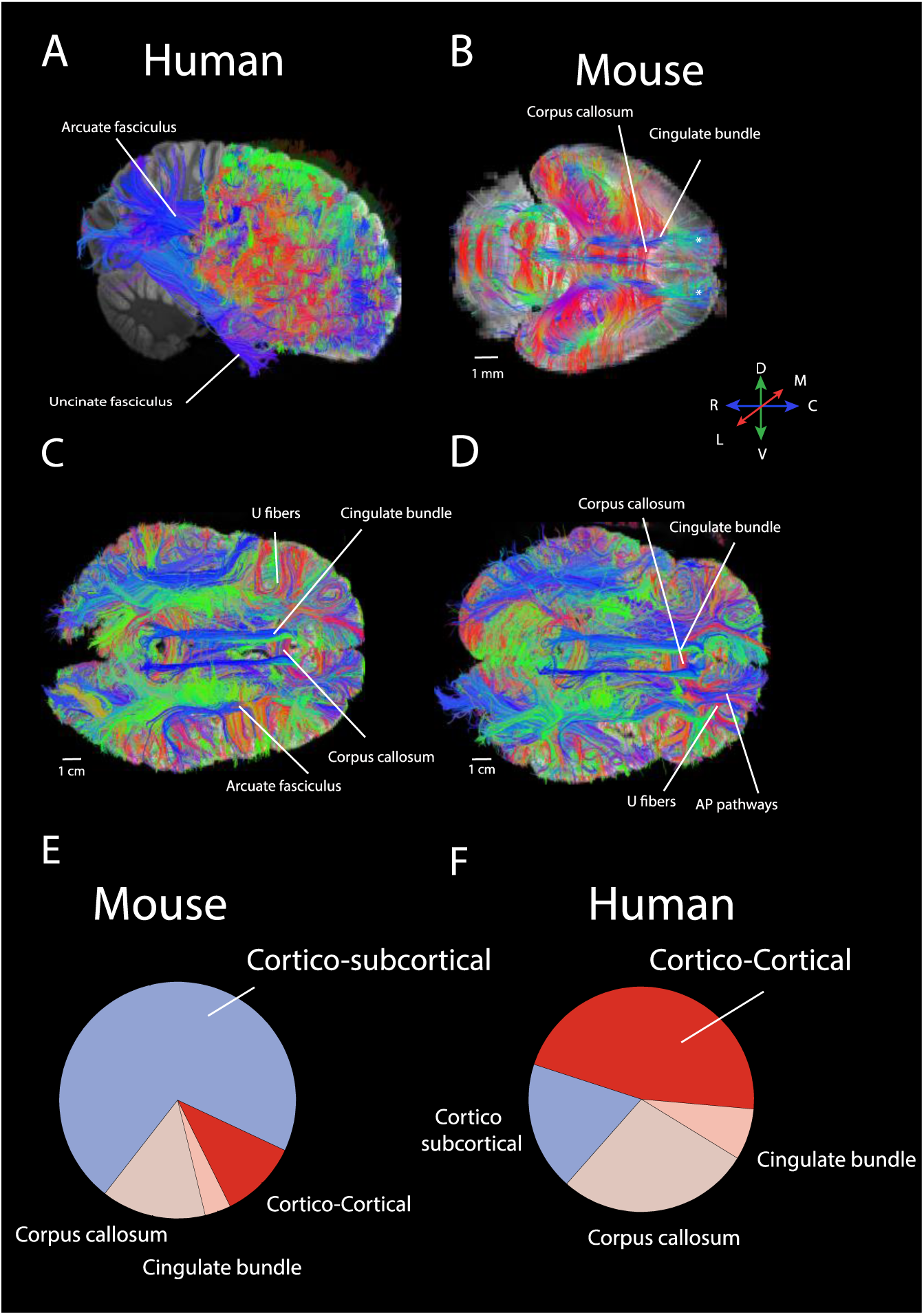
There are major differences in cortico-cortical pathways between humans (A) and mice (B) as assessed with high angular resolution diffusion MR tractography. Horizontal slice filters (C-D) and an ROI (A) set through the frontal cortex highlight a number of cortico-cortical pathways (e.g., arcuate fasciculus, uncinate fasciculus, U fibers, cingulate bundle) in humans. This is contrast with mice where only a select number of pathways are observed. Those are the corpus callosum, cingulate bundle, and the presumptive corticospinal tract (*). We quantified the relative number of identified pathways within the white matter of the frontal cortex. Pie charts show the relative number of identified pathways in the frontal cortex white matter of both species. The frontal cortex is preferentially composed of cortico-cortical pathways whereas the mice frontal cortex is preferentially composed of pathways connecting cortical and subcortical structures. Abbreviations: A: anterior; P: posterior; M: medial; L: lateral, D: dorsal; V: ventral.

To quantify these observed differences, we randomly placed ROIs through the frontal cortex white matter to capture fibers, and we classified observed pathways as belonging to the corpus callosum, cortico-subcortical, cingulate bundle, or other cortico-cortical pathways in both species in a total of 2 humans and five mice (Fig. 2-3, Fig. S2). We computed the relative number of identified pathway types (i.e., callosal, cingulate, other cortico-cortical, or cortico-subcortical) per individual (Fig. S2). Accordingly, the human frontal cortex white matter in humans is preferentially composed of cortico-cortical pathways relative to mice (Fig. 2E-F). A similar situation is observed when comparing a broader sample (Fig. 3). A t test on the relative number of cortico-cortical pathways in two humans and five mice show that humans possess significantly more cortico-cortical pathways 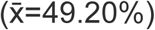 coursing through the white matter of the frontal cortex than mice (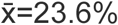; t=−3.2818, n=2 humans, n=5 mice, p<0.05; Fig. 3H). These findings demonstrate that increased expression of supragranular genes accompanies an expansion in long-range cortico-cortical projections emerging to or from the frontal cortex in humans.

**Figure 3.**
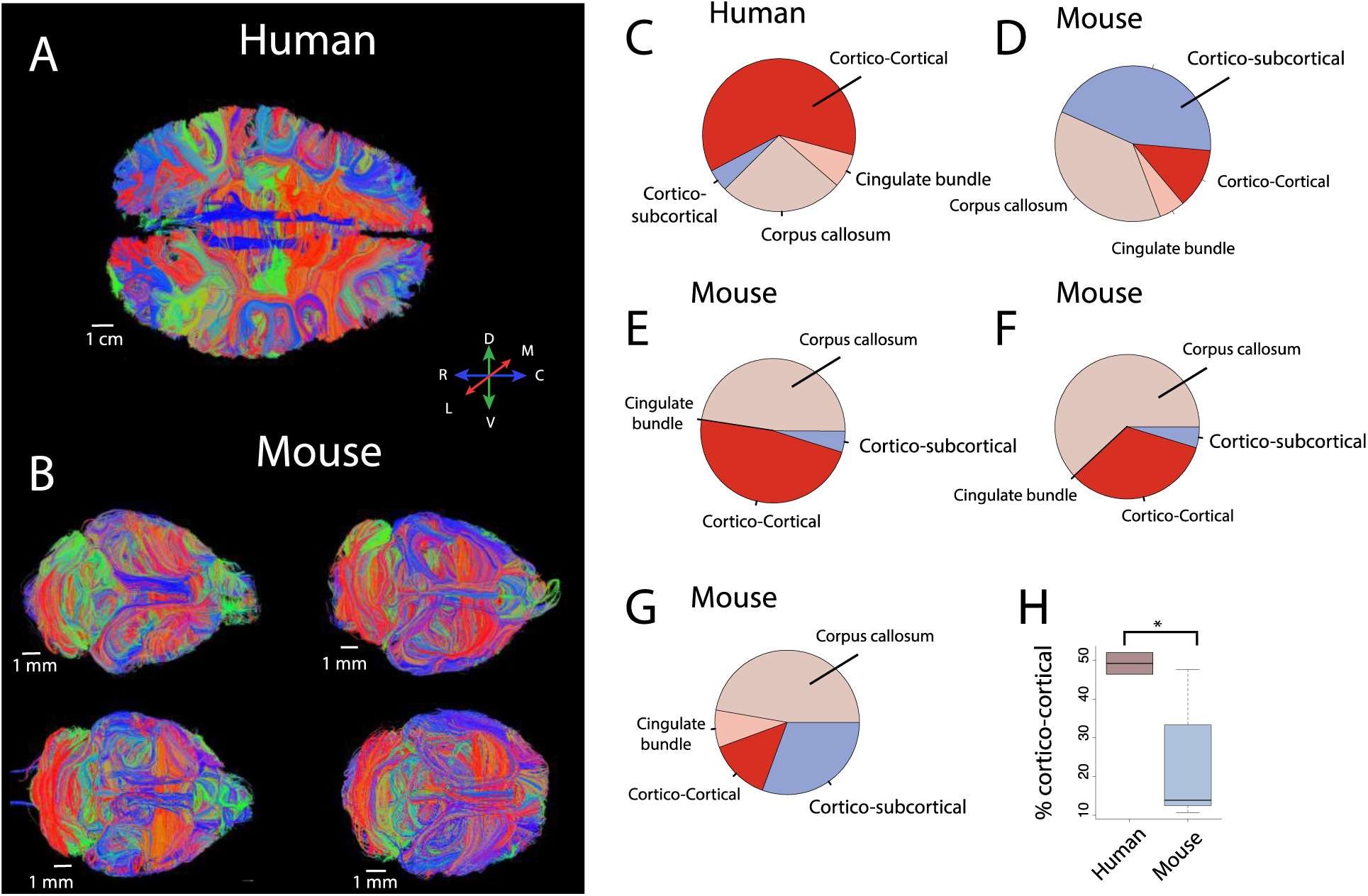
We compare the relative number of identified pathways coursing through the frontal cortex white matter of a 34 year old woman (A) made available via the Allen Brain atlas (Ding et al., 2016) and mice (B) with diffusion MR tractography. A comparative analysis of the relative number of identified pathways show that the frontal cortex white matter is preferentially composed of cortico-cortical pathway in humans (C) compared with mice (D-G). (H) The number of identified cortico-cortical pathways identified in the two humans are significantly increased relative to mice (n=5). Tracts coursing across the anterior to posterior axis are in blue. Tracts coursing lateral to medial are in red and tracts coursing across the dorsal-ventral axis are in yellow. Abbreviations: A: anterior; P: posterior; M: medial; L: lateral, D: dorsal; V: ventral.

### Developmental time course of supragranular-enriched genes in humans versus mouse frontal cortex

Comparative analyses of RNA sequencing data from the frontal cortex at successive ages can reveal differences in developmental trajectories gene transcription, and evolutionary modifications to biological pathways. We tested whether the developmental time course of supragranular-enriched genes are protracted in humans compared with mice with the use of RNA sequencing from bulk samples taken from frontal cortical regions of humans and mice at successive stages of development. We focus on supragranular enriched genes, which encode axon, voltage-gated sodium, potassium, and calcium channels, extracellular matrix, and synaptic-related genes, and compare developmental trajectories in both species after controlling for variation in developmental schedules. To compare the developmental trajectories of these genes, we mapped human age onto mouse age according to the translating time model (Workman et al., 2013). We rely on the timing of developmental transformations to find corresponding ages to map human age onto mouse age (Workman et al., 2013). Because the translating time model finds corresponding ages up to two years of age in humans and its equivalent across species, we extrapolated corresponding ages to span later times as we had done previously (Charvet and Finlay, 2018, Charvet et al., 2019). We find strong concordance between RNA sequencing data-sets (Lister et al., 2013, BrainSpan Atlas of the Developing Human Brain (2010); Fig. S3).

A cursory examination of some of these genes (e.g., *NEFH*, *VAMP1*, *SYT2*) shows that these genes peak or plateau in both species but that the expression of these genes steadily increase in humans for much longer than in mice once for variation in developmental schedules is controlled for (Fig. 4A-C), which suggest that the developmental profile of at least some of these supragranular-genes may be protracted in humans relative to mice. To confirm that the temporal pattern of at least some of these supragranular-enriched genes is indeed protracted in humans compared with mice, we compare the coefficient of variation of supragranular-enriched genes in humans and in mice at all and late stages of development (Fig. 4D). This analysis assumes that increased variation in one species relative to another would reflect evolutionary changes in developmental programs. We consider only expressed genes (FPKM>0.5 averaged across age ranges per species), which produced 18 supragranular-enriched genes for consideration. We fit a smooth spline through the expression of each gene versus age expressed in days after conception, and we interpolated corresponding ages across the two species. We retain fetal stages for mice, which correspond to earlier stages than that obtained for humans (Fig. 4A-C). We test for differences in the coefficient of variation in the expression of supragranular-enriched genes (Fig. 4D) and all expressed orthologous genes (Fig. 4E) across all or select age ranges in humans and mice. The coefficient of variation of supragranular-enriched genes is not significantly different between the two species when all ages are considered, suggesting there is not increased overall variation in the expression of supragranular-enriched genes in humans 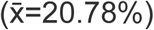 compared with mice (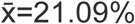; Fig. 4D; D=0.278; p=0.5, n=18). However, the coefficient of variation is significantly greater in humans than in mice once the examined age ranges span late time points (Fig. 4D). That is, the coefficient of variation is significantly increased in humans relative to mice once the examined age ranges span post-conception day (PCD) 50 and beyond (ks test: D=0.61, p<0.05, n=18; Fig. 4D) or PCD80 and beyond in mice (ks test, D= 0.55; p<0.05, n=18; Fig. 4D) and their equivalent ages in humans. In other words, humans possess prolonged variation in the expression of at least some supragranular-enriched genes relative to mice, which suggests a prolonged duration of development linked to the emergence and expansion of cortico-cortical pathways in humans.

**Figure 4.**
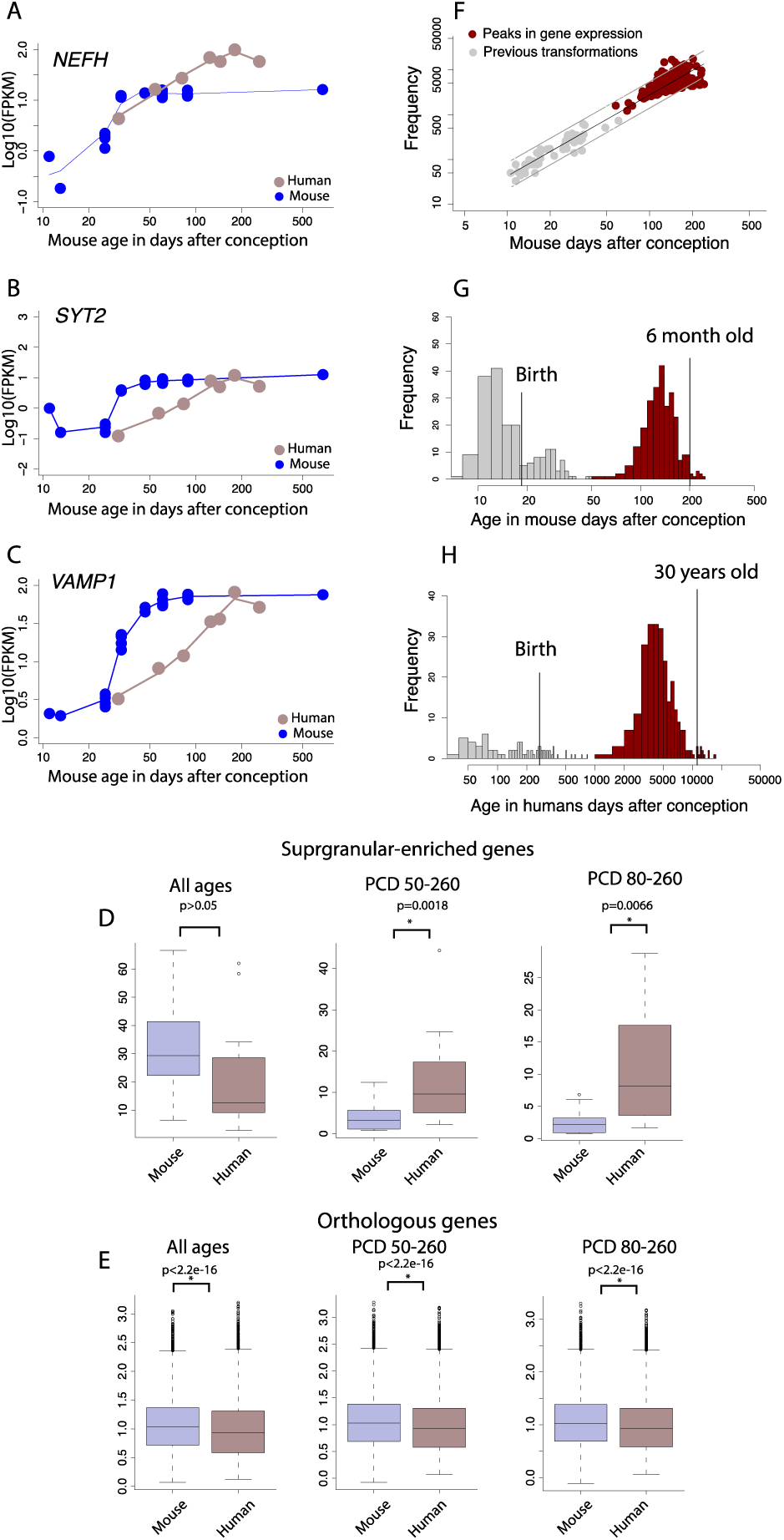
Developmental time course of (A) *NEFH*, (B) *SYT2*, and (C) *VAMP1* show a protracted developmental time course in humans relative to mice. (D) The coefficient of variation of supragranular - enriched genes across all examined age ranges are not significantly different between humans and mice. However, the coefficient of variation in the expression of supragranular-enriched genes are significantly increased in humans relative to mice if only later ages are considered, either from post - conception day (PCD) 50-260 or PCD 80-260 in mice and their equivalent in humans. (E) The increased coefficient of variation is not general to all expressed orthologous genes because the coefficient of variation is significantly less in humans compared with mice whether all age ranges are tested or whether only late stages are considered. (F) The timing in peak expression of genes is conserved enough to find corresponding ages across species. The timing of transformations and peaks in gene expression expressed in the log10 days after conception in mice is regressed against those in humans, and permit extending the timing of transformations to 30 years of age in humans (G) and 2 months of age in mice (H).

One caveat with this interpretation is that the expression of genes may simply be more variable in humans than in mice. The coefficient of variation of all expressed orthologous genes (n=10,815) is actually significantly less in humans it is in mice whether all examined age ranges are considered (D=0.09; p<2.2e-16; n=10,815), or late age ranges are examined (PCD and beyond: ks test, D= 0.08, p<0.05, n=10,815; PCD 80 and beyond, ks test D= 0.08, p<0.05, n=10,815; Fig. 4E). Therefore, the increased variation in supragranular-enriched genes in the human frontal cortex at late stages of development is not a product of overall increased variation in gene expression in the human frontal cortex, but is specific to supragranular-enriched genes. We also use temporal changes in gene expression to extrapolate corresponding ages from humans to mice during postnatal ages (Fig. 4F-H).

### Overall conservation in the timing of gene expression in humans and mice

We have extrapolated corresponding ages in humans and mice to span later time points. However, the translating time model only extends up to 2 years of age in humans and its equivalent in other species (Workman et al., 2013). It is, however, an open question as to whether it is actually possible to find corresponding ages at later stages of postnatal development. To ensure the validity of extrapolating the translating time model to later ages in both humans and in mice, we identify corresponding ages from either positive or negative peaks in the expression of orthologous genes from the frontal cortex of humans and mice (Fig. 4F-H). We fit a smooth spline though the log10 FPKM values in humans and in mice versus age expressed in days after conception, and extrapolate corresponding ages between humans and mice from around birth to 53 years of age, and their equivalent ages in mice. This resulted in 7 time points for comparison in the two species. We fit a quadratic model with the log-transformed normalized expression of each gene versus age in each species (Benjamini and Hochberg: BH FDR threshold: p<0.05) to determine the age in which a gene might peak in its expression in both species. We then only selected genes with a positive correlation between the two species, and corrected for multiple testing with an FDR threshold set to p<0.05. Such an analysis yielded 261 genes, which significantly correlate, and peak either positively or negatively in both species. We compare corresponding ages extracted from temporal changes in gene expression from the frontal cortex with previously collected transformations used to find corresponding ages across species (n=59; Charvet and Finlay, 2018). We fit a linear model with the logged values of developmental event timing from previously collected transformations and peaks in gene expression expressed in age in days after conception in mice and humans. This model accounts for 94.31% of the variance (y=1.80x-0.17, df=318, p<0.05; Fig. 4F), and extend the age ranges to 30 years of age in humans (Fig. 4G) and 6 months of age in mice (Fig 4H). Integrating the age of peaks in gene expression with anatomical transformations is an effective strategy to find corresponding ages at later stages of development in humans and in mice.

### Growth of frontal cortex white matter in humans and mice

We compare the timetable of long-range cortical pathways emerging to and from the frontal cortex in humans as in mice by comparing the growth of frontal cortex white matter left hemisphere across the two species to assess whether the development of long-range projections emerging to or from the frontal cortex is protracted in humans relative to the timing of other transformations (Fig. 5A-C). In both species, the frontal cortex white matter grows postnatally but eventually ceases to grow. Human age is mapped onto mouse age according to the translating time model (Fig. 5A, C). We used a non-linear model to identify (easynls, model=3) to identify when the frontal cortex white matter ceases to grow across 83 humans and mice and averaged frontal cortex white matter volumes per time point. The frontal cortex white matter ceases to grow on post-conception day 47 in mice (y=0.03+0.015x(x-47.22); R^2^=0.70; p(coefficient a)=0.61, p (coefficient b)<0.05; p(coefficient c)<0.05; n=12) and around 5.5 years of age in humans. In humans, 5.5 years of age is equivalent to post-conception day 88 in mice (y=4.83+0.01x(x-88.43), R^2^=0.81, p(coefficient a)<0.05, p(coefficient b)<0.05, p(coefficient c)<0.05, n=35). The age of frontal cortex white matter cessation occurs later than expected in humans once variation in developmental schedules are controlled for (Fig. 5A, C). Events that occur between post-conception day 66-80 in mice occur on 1.3-1.5 years of age (PCD 735-833; n=2) in humans (Workman et al., 2013). The timing of frontal cortex white matter growth cessation falls outside the 95% CI of these data (PCD 828). The timing of frontal cortex white matter cessation of humans falls outside the 95% prediction intervals logged values of developmental event timing from transformations and peaks in gene expression expressed in age in days after conception in mice and humans (Fig. 5D). Therefore, the growth of the frontal cortex white matter, which houses projections coursing within the white matter of the frontal cortex, is protracted in humans relative to the timing of other transformations.

**Figure 5.**
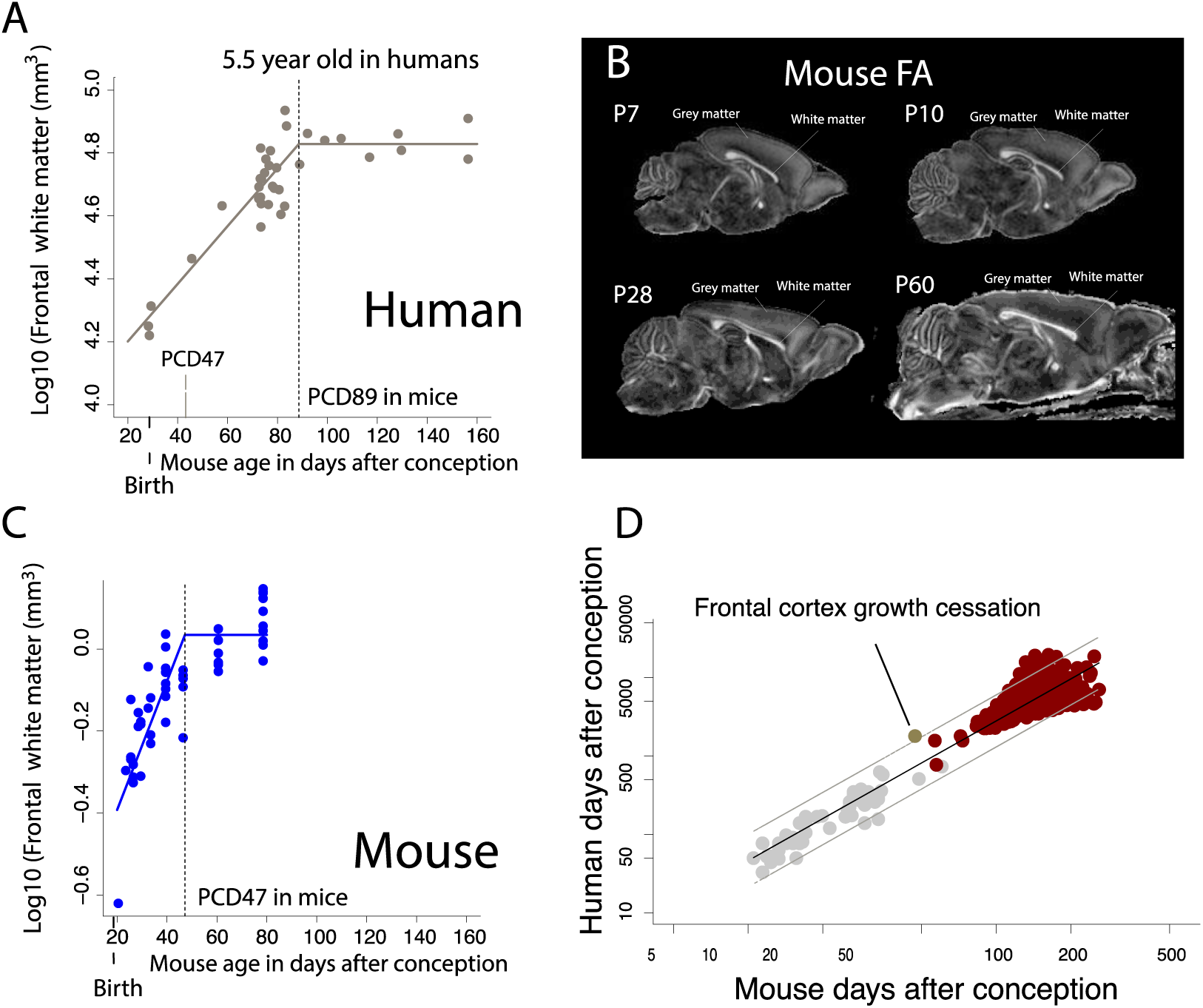
We measure the growth of the frontal cortex white matter in humans (A) and in mice (B) from structural MR scans in humans or FA images from diffusion weighted images in mice (C). In mice, the frontal cortex white matter was visualized from FA images of diffusion weighted scans at different ages. (A) The frontal cortex white matter growth in humans is mapped onto mouse age according to the translating time model (Workman et al., 2013). After controlling for variation in developmental schedules across the two species, the frontal cortex white matter growth is extended in humans (A) compared with mice (B). According to the non-linear regression used to capture frontal cortex white matter cessation, the frontal cortex white matter ceases to grow at about 5.5 years of age in humans, which corresponds to post-conception day 89 in mice. In contrast the frontal cortex white matter ceases to grow relatively earlier in mice (post-conception day 47). Vertical dashed lines represent the timing of frontal cortex white matter growth cessation in both species. (D) To further ensure that the frontal cortex white matter is indeed protracted in humans compared with the timing of other transformations, we compare the timing of frontal cortex white matter cessation growth in humans and mice relative to the timing of transformations assessed from prior work as well as from peaks in gene expression. To that end, we regress the log10 values of the timing of developmental transformations in humans and in mice and generate 95% prediction intervals from these data. The age of frontal cortex white matter cessation falls outside the 95% prediction intervals from these data. The growth of the frontal cortex white matter is protracted in humans relative to the timing of other transformations.

## Discussion

The present study focuses on comparative analyses of pathways from the frontal cortex from diffusion MR imaging and supragranular-enriched genes in order to identify evolutionary changes in connectivity patterns in adulthood and in development. Our findings demonstrate that integrating RNA sequencing data with neuroimaging phenotypes is an effective approach to identify evolutionary changes in connections in adulthood as well as in development. We also demonstrate that considering temporal changes in gene expression is an effective strategy to find corresponding ages between humans and mice during postnatal development and in adulthood.

Integrating comparative analyses from diffusion MR tractography with supragranular-enriched genes converge on the observation that the human frontal cortex is preferentially composed of layer III neurons, concomitant with increased cross-cortically projections through the white matter of the frontal cortex in humans compared with mice. We use diffusion MR tractography and find an expansion in cortico-cortical pathways in the frontal cortex of humans, which fall in line with predictions from our comparative analyses of supragranular-enriched genes. We further demonstrate that the developmental trajectories of supragranular-enriched genes and growth of projections emerging to and from the frontal cortex are both protracted in their developmental profile in humans. Thus, the expansion of layer III neurons and concomitant expansion of projections coursing within the white matter of the frontal cortex in humans is achieved by extending the duration of their development. We have yet to identify the extent to which such modifications are unique to humans, shared among primates, and how such modifications to developmental programs vary across mammals.

Several hypotheses have been proposed to account for variation in connectivity patterns across species, but were not explicitly tested due to the lack of methods available to quantify variation in connectivity patterns across species in adulthood and in development (Innocenti, 1995; Striedter, 2005). Tract-tracers, diffusion MR tractography, and carbocyanine dyes on their own suffer from a number of technical limitations (Nudo and Masterton, 1989, 1990; Chen et al., 2006; Reveley et al., 2015; Heilingoetter and Jensen, 2016). Diffusion MR tractography identifies pathways across the entire brain, but suffers from a number of false positives, especially in *in vivo* (Thomas et al., 2014; Reveley et al., 2015). As a result, hypotheses focused on the evolution and development have largely gone unexplored. Our work integrates neuroimaging and molecular level phenotypes to identify deviations in connections, and their underlying developmental programs. We demonstrate that such an approach is an effective strategy to address evolutionary modifications to developmental programs leading to connectivity patterns in the human frontal cortex.

As is evident in the present manuscript, the establishment of connections extends post-nally in both humans and mice. Comparing developmental programs underlying variation in connections across species relies on identifying corresponding time points across species. We here use temporal trajectories in gene transcription from the frontal cortex in both species to find corresponding time points up to 30 years of age in humans and their equivalent in mice. Identifying sources of conservation and variation in developmental programs during postnatal ages is an essential enterprise as it will permit better extrapolating findings from model organisms to humans. This is because researchers often use model organisms during postnatal development and in adulthood to understand basic developmental processes and dysfunctions that occur in humans (Charvet and Finlay, 2018). Extrapolated ages from the translating time model fall in line with our results extracted from peaks in gene expression from the frontal cortex of both species (Workman et al., 2013). Controlling for variation in developmental schedules reveals that the maturation of pathways emerging to or from the frontal cortex is protracted relative to the timing of other transformations in humans relative to mice.

Modifications to the timing of developmental programs have repeatedly been demonstrated as an important source of variation in brain structure and function across vertebrates. For instance, the disproportionate expansion of the cortex and cortical neuron numbers in human and non-human primates is achieved by extending the duration of cortical neurogenesis relative to the timing of most other transformations (Charvet et al., 2017, 2019b). The extension in the duration of cortical neurogenesis accounts for the amplification of upper layer (i.e., layer II-IV) neurons, which participate in local and long-range cortico-cortical projections (Cahalane et al., 2014; Charvet et al., 2017a, 2019). We demonstrate yet another important developmental change in the timing of biological pathways (i.e., heterochrony) in the human lineage. That is, extending the development of projections accounts for their expansion within the human frontal cortex, and the over-representation of frontal cortex layer III neurons in humans. We have yet to identify whether such variation is specific to the frontal cortex or general to the isocortex. The extension in the duration of frontal cortex development might permit increased time to learn from conspecifics during a prolonged duration of infancy and adolescence in humans. This work will provide an opportunity to link evolutionary changes in the timing of behavioral and cortico-cortical pathway development across species in the future.

The framework as articulated here may be applied to resolve ongoing debates focused on the evolution of the human brain. One example is whether the human frontal cortex, and in particular the prefrontal cortex white matter, is unusually large compared with apes (Donahue et al., 2018). The lack of consensus across studies suggests the need to move beyond gross measures of white matter, and to focus on integrating additional measures of connectivity at multiple scales of organization across species. The set of supragranular-enriched genes coupled with diffusion MR tractography provides an opportunity with which to investigate the evolution of long-range cortico-cortical pathways more closely in the human lineage, and to resolve whether the expansion in long-range cortico-cortical pathways within the white matter of the frontal cortex is unique to humans or shared among primates.

## Materials and Methods

### Comparative analyses of layer III of adult humans and mice

We performed a number of analyses to confirm the expansion of layer III neurons and cortico-cortical pathways in the human frontal cortex compared with mice (Fig. 1). Those include comparative analyses from diffusion MR tractography (Fig. 2-3), as well as gene expression from bulk and single cells to confirm that humans do indeed possess an expansion of long-range cortico-cortical projections in the frontal cortex compared with mice (Fig. 4; Zeng et al., 2012; Charvet et al., 2017a, 2019a). We compared 1) the relative thickness of layer III across the frontal cortex of humans and mice with use of *CALB1* expression, 2) the relative number of layer III neurons in humans and in mice from a previously published single cell methylome sequencing study (Luo et al., 2017), and the relative expression of supragranular-enriched genes across the layers of the human and mouse frontal cortex. We also compared cortico-cortical pathways with the use of diffusion MR tractography to test whether cortico-cortical pathways are expanded in humans compared with mice.

We first measured the relative thickness of supragranular layers of adult human and mice frontal cortices. Supragranular layers were defined by the expression of *CALB1* in humans and mice because *CALB1* is a marker across supragranular layers. Thickness measurements were taken from *CALB1* in situ hybridization (ISH) images of humans from 28 to 31 years of age (n=3) and mice at post-natal day 56 (n=4) made available as part of the Allen brain atlas (2010 Allen Institute for Brain Science; Fig. 1). A grid was placed on sections of either human or mouse frontal cortices, and we randomly selected sites to measure the thickness of layer III from *CALB1* ISH sections across the depth of the cortex. We used adjacent Nissl stains to measure the thickness of layers I-VI. Layer VI was bounded by the white matter. The relative thickness of *CALB1* is computed as layer III versus layer I-VI thickness, and we performed a t test on these data. We applied the same approach to measure the relative thickness of layer III from Nissl-stained sections (Fig. S1).

We measured the intensity of supragranular-enriched expression of some genes across upper (i.e., layers II–IV) versus lower (i.e., layers V–VI) layers in the frontal cortex of humans and mice (i.e., *CRYM*, SYT2, *VAMP1*, *NEFH*. Previously, supragranular-enriched genes were shown to be preferentially expressed in supragranular layers of the human primary visual cortex relative to mice, but such differences were not demonstrated in the frontal cortex. To quantify differences in mRNA expression across layers of the frontal cortex, we downloaded in situ hybridization images for *CRYM*, *SYT2*, and *VAMP1* made available by the Allen Brain Atlas. We placed a rectangular grid through at least two sections per region per individual with 4-6 sections per individual as we had done previously to quantify the relative intensity of these genes across upper and lower layers (Charvet et al., 2019a). Randomly selected frames were aligned along the cortical surface. Frame widths were 500 and 1,000 μm in mice and humans, respectively. The height of frames varied with the thickness of upper and lower layers. These analyses were performed in Image J. We used cytoarchitecture from adjacent Nissl-stained section in humans, and in mice, as well as atlases to define cortical areas, as well as upper and lower layers. Upper layers were distinguished from lower layers based on a small cell-dense layer IV evident from Nissl-stained sections. This approach is similar to that used previously to quantify densities in *NEFH* expression across upper and lower layers, which showed increased *NEFH* expression in upper layers in humans relative to mice (Charvet et al. 2015, 2019a). We also include these comparisons of *NEFH* expression, which were obtained in a manner identical to that described previously (Charvet et al., 2019a).

### Comparative analyses of diffusion MR tractography in humans and mice in adulthood

We compare the relative number of pathways emerging to and from the frontal cortex of humans and mice (Fig. 2-3). A total of two post-mortem human and 5 mouse brains were placed in a perfluoropolyether (Fomblin; Solvay, Brussels, Belgium) solution (invisible in MRI). The human brains were scanned with a 3T clinical Siemens MRI scanner (Erlangen, Germany) at the A. A. Martinos Center for Biomedical Imaging. We selected a steady-state free precession sequence with a repetition time (TR) of approximately 38 ms and an echo time (TE) of approximately 25 ms for diffusion acquisition (McNab et al., 2009). Forty-four diffusion-weighted images (b=1,000 s/mm^2^) and four non diffusion-weighted images were acquired. The b-value with the steady-state free precession sequence is T1-dependent. The other post-mortem human brain is made available via the Allen Brain Institute, and details of procedures have been described previously (Ding et al., 2016).

The mouse brains were scanned with a 4.7 T Bruker Biospec MRI system equipped with a high-performance gradient and a radio-frequency coil that best fit the small brains at the A. A. Martinos Center for Biomedical Imaging. A three-dimensional diffusion-weighted spin-echo echo planar imaging (SE-EPI) sequence was used with a TR of approximately 1,000 ms and TE of 40 ms to image the mouse brain. Sixty diffusion-weighted measurements (b=8,000 s/mm^2^) and one non-diffusion-weighted measurement (b=0) were acquired. These brains were used in previous studies, and details of scanning procedures have been described in more detail (Charvet et al., 2017a, 2019b).

We used high angular resolution diffusion imaging (HARDI) to detect pathways in human and mouse brains to detect fibers coursing in multiple directions within a voxel. Diffusion MRI data were processed with Diffusion Toolkit (www.trackvis.org). All orientation distribution functions (ODFs) were normalized by the maximum ODF length within each voxel, and calculated fractional anisotropy (FA) from orientation vectors by fitting the data to the usual tensor model (e.g. Takahashi et al., 2010). We used a FACT (fiber assessment by continuous tracking) algorithm to reconstruct pathways. No fractional anisotropy threshold was applied in reconstructing tracts, which is an approach consistent with that of other studies (e.g. Schmahmann et al., 2007; Takahashi et al., 2010, 2011; Kanamaru et al., 2017). We used TrackVis (http://trackvis.org) to visualize pathways. We set a horizontal slice filter (3mm thick) through the brain of the mouse at the level of the frontal cortex white matter to capture pathways for display purposes in Figure 2. A min fiber length=5mm; max fiber length=20 mm) coursing to and from the frontal cortex was set for visualization purposes in mice (Fig. 2). We set an ROI through the white matter of the frontal cortex in humans to capture pathways coursing to and from the frontal cortex. No minimum length fiber was applied to the human brain.

To quantify differences in cortico-cortical pathways between the two species, we randomly selected an ROI consisting of a voxel in the white matter of the frontal cortex of the human and the mouse, and we identified whether the fibers coursing through the randomly selected ROI consist of the corpus callosum, the cingulate bundle, other cortico-cortical pathways, or pathways connecting cortical and subcortical structures (Fig. S2). If a pathway was observed coursing through the dorsal midline, it was considered to be callosal. If the tractography neither clearly targeted subcortical, or cortical structures, we did not classify the pathway. Pathways connecting cortical and lateral limbic structures were considered to be cortico-subcortical pathways. Other cortico-cortical pathways consisted of U fibers as well as long-range projection pathways (e.g., arcuate fasciculus). We randomly selected 6-13 sites and 12 to 31 sites across serial coronal planes through the unilateral frontal cortex in mice and humans, respectively. More ROIs were selected through the human brain than in mice because the frontal cortex of humans is larger than it is in mice. We then computed the relative number of identified pathway types (i.e., callosal, cingulate, other cortico-cortical, or cortico-subcortical) per individual, and performed a t test on the relative number of observed other cortico-cortical pathways in humans versus mice.

### Developmental trajectories in gene expression in human and mouse frontal cortex

We use a previously published RNA sequencing dataset extracted from the human (mid-frontal gyrus; n=7; around birth to 55 years of age) and mouse frontal cortex with ages ranging from embryonic day 11 to 22 months of age (frontal cortex; n=21; Lister et al., 2013). Expression values are in fragments per kilobase million (FPKM; GEO accession number: GSE47966; Lister et al., 2013). Gene expression was quantified as the number of reads aligned to the gene exons divided by the length of the gene exon. To correct for variation in library size across samples, the mean number of reads across all collected samples was divided by the number of reads from the sample of interest. Only reads in coding regions, which could be lifted over (with the liftOver tool) across species were considered. Replicates and number of mapped reads are shown in Fig. S4. All statistical analyses were performed with the programming language R.

We filtered the dataset to consider orthologous expressed genes with an “OK” status. If a gene has several copies, we selected the gene with the greatest expression. We identified orthologous genes as defined by the mouse genome database (Smith et al., 2018). Only genes with an average expression threshold above 0.5 FPKM across the examined ages for each species were considered (mice: postnatal day: 3 to 8 months after birth; humans: 35 days after birth to 52 years of age). These filtering steps resulted in 18 out of 19 supragranular genes for consideration. Of these supragranular-enriched genes, *NEFH*, *VAMP1*, and *SYT2* reach a plateau of expression in both species. Notably, *NEFH* (a marker of long-range projecting neurons), *SYT2* expression (synaptic-related), and *VAMP1* appear to plateau in their expression later in humans than in mice once variation in developmental schedules between these two species are controlled for (Fig. 4; Charvet et al., 2019b). Given this differential pattern of gene expression between these two species, we test whether the coefficient of variation in the expression of supragranular-enriched genes is greater at late stages of development in humans compared with mice.

We assume that genes that vary with age in one species but not the other reflect deviations in developmental program between the two species. To that end, we fitted a smooth spline through the log10 of FPKM versus the log10 of age expressed in days after conception for each gene in each species. We then extrapolated corresponding ages across the two species to compare equivalent developmental time points, but we retained fetal stages as it appeared that, at least, some of these gene vary during fetal development in mice. Across all analyses, we added a value of 1 to all logged10 FPKM values to consider genes that may not be expressed at a particular time point. We then performed a t test on the coefficient of variation of supragranular-enriched genes and all orthologous genes in humans and mice across different age ranges to identify whether the variation in the expression of supragranular genes is significantly increased at late stages of development in humans compared with mice (Fig. 4). We repeated the same analyses across all orthologous genes to ensure that gene expression is not simply more variable in humans than in mice. To ensure that the temporal patterns of at least some of these supragranular-enriched genes are consistent with other RNA data-sets, we compare the temporal patterns of *NEFH* expression and *VAMP1* from the frontal cortex of humans and mice with that from the Allen Brain Atlas (Supplementary Figure 1; Lein et al., 2007; Miller et al., 2014; Hawrylycz, M.J. et al. 2012; Allen Developing Mouse Brain Atlas, 2008; BrainSpan Atlas of the Developing Human Brain, 2010).

### Temporal changes in gene expression to find corresponding ages at late stages of development

We next test whether temporal changes in gene expression in the frontal cortex are conserved enough to find corresponding ages across species. To that end, we selected genes with an average FPKM expression value above 0.5. To find corresponding ages at late stages of development, we fit a smooth spline through the logged values of FPKM versus the log-transformed values for age in mice to predict FPKM values at ages that are equivalent to those in humans. Because fitting a smooth spline should reduce variance around the mean, we fit a smooth spline when regressing logged values of FPKM versus the log-transformed values for age in humans as well. These smoothing steps resulted in a total sample of 7 samples of humans and in mice to be tested for non-linear associations.

For each species, we tested which genes exhibit a significant peak in their expression with age with easynls package in R (model 2). The age in which a plateau of expression was observed was constrained to span the same age ranges in both species. We then corrected for multiple testing with a false discovery rate (FDR) BH threshold p value set to 0.05. We filtered the number of associations further to only include expressed genes that significantly correlated in both species. To that end, we correlated (cor) the log10 based FPKM values in humans (n=7) and in mice (n=7) matched for age. We then selected genes that correlate positively and significantly between the two species, and corrected for multiple testing with an FDR threshold set to p<0.05. This resulted in 261 genes with a significant peak in expression in both species. We then test whether these data can be used to extrapolate corresponding ages to later time points in both species. To that end, we include prior work capturing the timing of developmental transformations in humans and mice (Charvet and Finlay, 2018). To increase the number of developmental transformations for consideration in mice, we extrapolated corresponding ages between rats and mice by fitting a linear regression through the log-transformed values in the timing of developmental transformations for rats versus those in mice (y=1.05x-0.11, R^2^=0.96, df=137). We then extrapolated corresponding ages from rats to mice, which resulted in 58 developmental transformations in mice (Fig. 4F).

### Developmental trajectories of frontal cortex white matter growth in humans and mice

To assess whether there are deviations in the development of white matter pathways between humans and mice, we measured the volume of the frontal cortex white matter in the left hemisphere at successive stages of development in 36 humans (gestational week 36 to 18 years old) and 47 mice (P2 to P60). We used *in vivo* and *ex vivo* structural and diffusion MRI scans to capture the growth of the frontal cortex white matter in humans and in mice, respectively (Shi et al., 2011; Chuang et al., 2011; Khan et al., 2018a,b; Richardson et al., 2018). In humans as in mice, we used atlases to demarcate the white matter of the frontal cortex from the remaining cortical areas (Ding et al., 2016). The anterior cingulate cortex was included as part of the frontal cortex if the frontal cortex was observed in the same plane of view. In mice, the caudal or posterior boundary between the frontal cortex and remaining cortex was bounded by the presence of the hippocampus from coronal planes. We used atlases (Paxinos and Franklin, 2019) to define the lateral boundary between the frontal cortex and somatosensory cortex, which was most challenging to delineate in mice. We used FA images from diffusion weighted images in mice because FA permitted demarcating the frontal cortex white matter at successive stages of development and differences between grey and white matter were clearly observed (Fig. 5B). In humans, we selected structural MR scans because structural MR scans yielded better resolution than FA images. A minimum of 5 serial planes were used to reconstruct the volume of the frontal cortex in humans and in mice. We multiplied the area by the section spacing to compare volumes of frontal cortex white matter over the course of development and across species. We compare the timing of frontal cortex white matter growth cessation between the two species to test whether the frontal cortex white matter grows for longer than expected in humans compared with mice. In mice as in humans, we averaged frontal cortex white matter volumes for each age prior to testing for a linear plateau with easynls (model-3) implemented with the programming language R. We fit a non-linear regression with the log10-transformed values for the frontal cortex versus age expressed in days after conception. Age in humans was mapped onto mouse age according to the translating time model (Workman et al., 2013; http://translatingtime.org). Because the observed age ranges span later time points than those predicted by the translating time model, we extrapolated corresponding ages to later ages in humans and mice as we have done previously (Charvet et al., 2017, 2019ab).

## Acknowledgements

We thank Dr. Melissa Harrington for her support at Delaware State University. A developmental series of mouse FA scans were obtained as a courtesy of Dr. Mori (lbam.med.jhmi) at Johns Hopkins University available at http://cmrm.med.jhmi.edu. In situ hybridization data and associated images from mice and humans were taken from the Allen Institute Website and the Brainspan atlas of the developing human brain. These data are available at http://developingmouse.brain-map.org and at http://www.brainspan.org, which are supported by the National Institute of Health Contract HHSN-271-2008-00 047-C to the Allen Institute for Brain Science. The opinions in this article are not necessarily those of the NIH. This work was supported by National Institute of General Medical Sciences (NIGMS) grant (5P20GM103653), and National Institute of Neurological Disorders and Stroke (NINDS) grant (R25NS09537) for research at Delaware State University. This work was also supported by the Eunice Shriver Kennedy National Institute of Child Health and Development (NICHD) (R01HD078561, R21HD069001) (E.T.).

---

**Figure S1.**
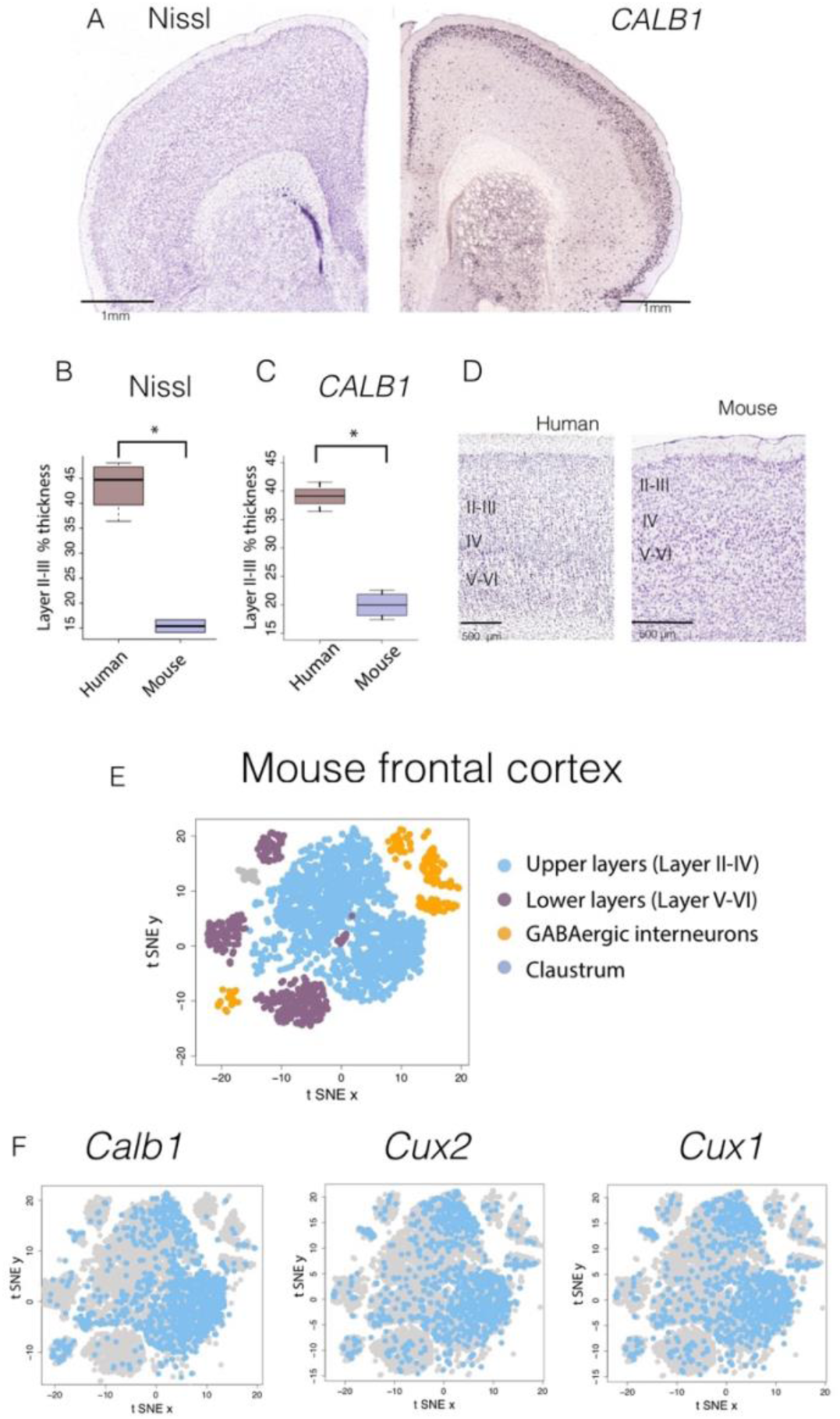
*CALB1* is an excellent marker for layer II-III neurons in the frontal cortex. (A) Nissl stains and *Calb1* expression from coronal planes of the mouse frontal cortex show that *Calb1* expression is constrained to upper layers of the mouse cortex. (B) The relative thickness of layer II-III as assessed with (B) Nissl stains and (C) *CALB1* yield highly similar results with the relative thickness of layer II-III significantly increased in humans compared with mice. (D) Close-up views through layer II-IIII in humans and mice show that the cortex is preferentially composed of layer II-III. (E) We use tSNE plots of RNA expression from single cells extracted from the frontal cortex of mice identifies clusters of neuronal populations to ensure that *Calb1* is a good marker for layer II-III neurons in mice. Populations as identified as layer II-IV, layer V-VI, GABAergic interneurons, and claustrum cell populations are color-coded. (F) The expression *Calb1* is largely restricted to upper layers. Moreover, the pattern of *Calb1* expression resembles those of well known layer II-III makers such as *Cux1* and *Cux2*. Data and identified clusters were from generated by Saunders et al. 2018. A gene was considered to be expressed if its expression value was greater than 0.

**Figure S2.**
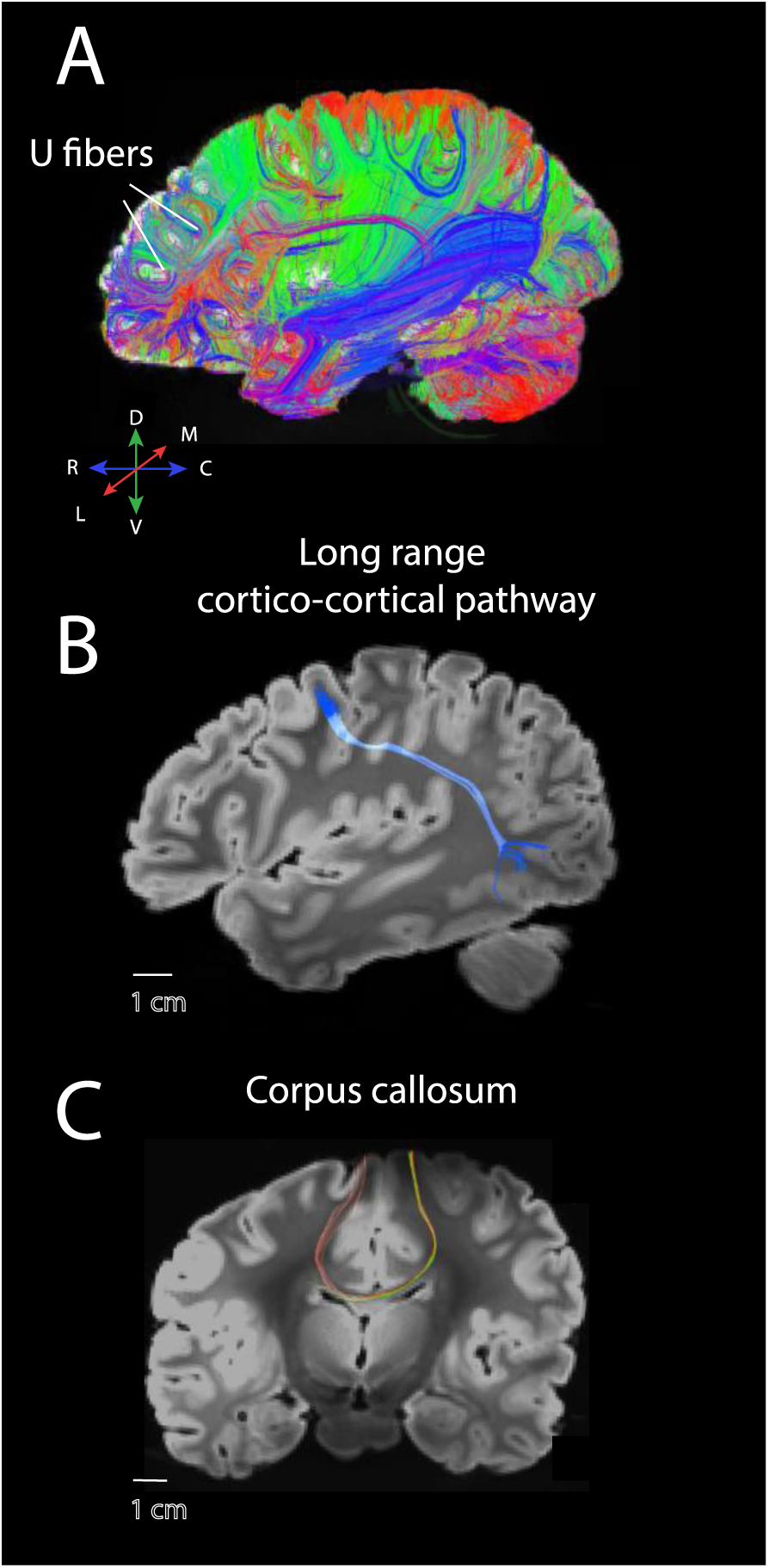
Examples of pathways observed coursing through the white matter of the frontal cortex. (A) Sagittal slices through the human brain show a number of fibers coursing across the anterior to posterior and medial to lateral direction within the frontal cortex white matter, including U fibers coursing through the frontal cortex white matter. Randomly selected ROIs through the white matter of the frontal cortex revealed different classes of cross-cortically projecting fibers coursing to or from the frontal cortex. Those include long-range cross-cortically projecting pathways coursing across the anterior to posterior direction (B), as well as the corpus callosum (C). Abbreviations: A: anterior; P: posterior; M: medial; L: lateral, D: dorsal; V: ventral.

**Figure S3.**
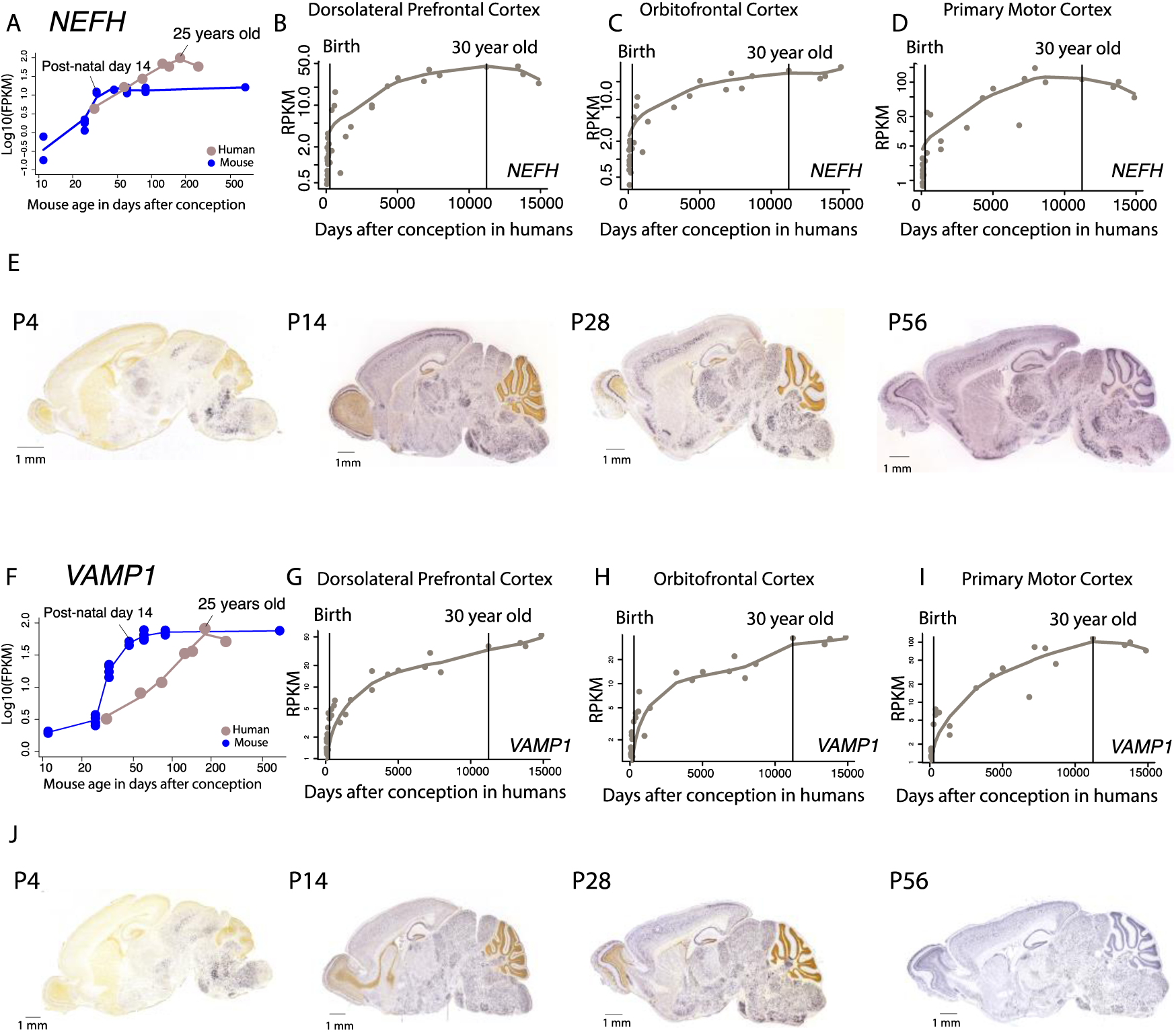
We find strong concordance in the temporal pattern of *NEFH* and *VAMP1* across data-sets. We compare the time course in expression of these genes from Lister et al., 2013 with that made available from the Allen Brain Institute (BrainSpan Atlas of the Developing Human Brain (2010)). (A) In humans, *NEFH* expression in the frontal cortex increases up to about 25 years. Similarly, in humans, the expression of *NEFH* expression from the dorsolateral prefrontal cortex (B), the orbitofrontal cortex (C), and the primary motor cortex (D) increase up to approximately 30 years of age. In mice, in situs show that *Nefh* expression increases substantially between post-natal day (P) 4 to P14 in the mouse cortex to reach stable levels thereafter. These observations are consistent with those from Lister et al,, 2013. A similar situation is observed for *VAMP1* where (F) *VAMP1* steadily increases postnatally up to about 25 years of age in humans. RNA sequencing data from the Allen Brain Institute shows that *VAMP1* increases up to around 30 years of age and somewhat thereafter in the dorsolateral prefrontal cortex (G), the orbitofrontal cortex (H), and the primary motor cortex (I) in humans. In mice, sagittal sections of in situs show that *VAMP1* increases steadily between P4 to P14 to reach relatively stable levels thereafter. These data are highly consistent with those of Lister et al., 2013 which show that *VAMP1* steadily increases up to about 25 years of age but that *VAMP1* expression remains relatively invariant after post-natal day 14 in mice.

**Figure S4.**
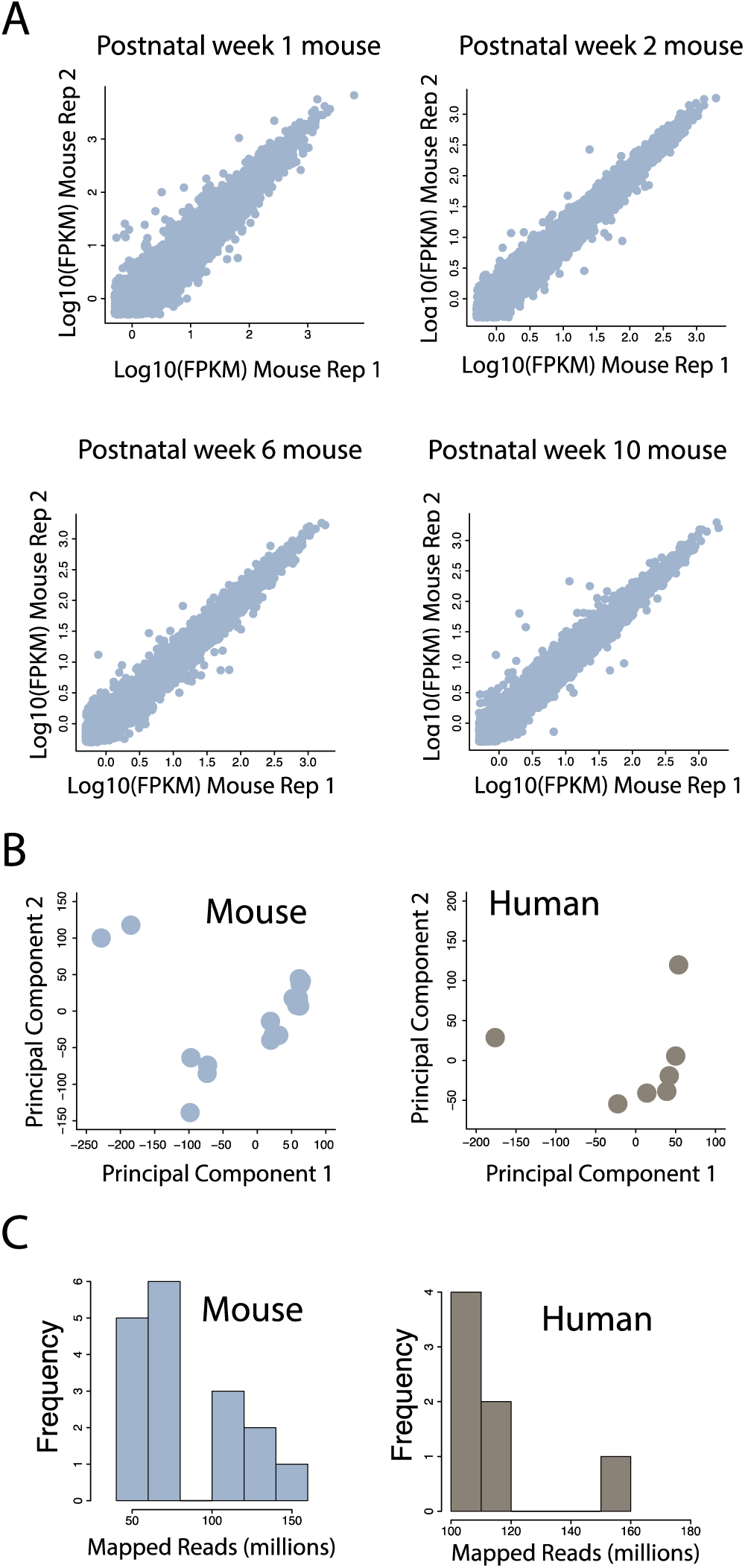
(A) Example of RNA sequencing replicates through the frontal cortex of mice at different ages. Minimum expression of 0.5 FPKM was selected across replicates. (B) Principal component analysis on the individuals extracted from frontal cortical areas of humans (n=7) and mice (n=21). Only expressed genes were selected, and no outliers were removed based on the principal component analyses. (D) Histogram of RNA sequencing reads mapped to the genome extracted from the frontal cortex of humans (n=7) and mice (n=21). Mapped reads ranged between ~54 and ~143 million in mice, and ~106 to 155 million in humans.

